# BMP-Dependent Mobilization of Fatty Acid Metabolism Promotes *Caenorhabditis elegans* Survival on a Bacterial Pathogen

**DOI:** 10.1101/2025.03.13.643118

**Authors:** Katerina K. Yamamoto, Margaret Wan, Rijul S. Penkar, Cathy Savage-Dunn

## Abstract

The Bone Morphogenetic Proteins (BMPs) are secreted peptide ligands of the Transforming Growth Factor beta (TGF-β) family, initially identified for their roles in development and differentiation across animal species. They are now increasingly recognized for their roles in physiology and infectious disease. In the nematode *Caenorhabditis elegans*, the BMP ligand DBL-1 controls fat metabolism and immune response, in addition to its roles in body size regulation and development. DBL-1 regulates classical aspects of innate immunity, including the induction of anti-microbial peptides. We theorized that BMP-dependent regulation of fat metabolism could also promote resilience against microbial pathogens. We found that exposure to a bacterial pathogen alters total fat stores, lipid droplet dynamics, and lipid metabolism gene expression in a BMP-dependent manner. We further showed that fatty acid desaturation plays a major role in survival on a bacterial pathogen, while fatty acid β-oxidation plays a more minor role. We conclude that *C. elegans* mobilizes fatty acid metabolism in response to pathogen exposure to promote survival. Our investigation provides a framework to study potential metabolic interventions that could support therapeutics that are complementary to antibiotic strategies.

## Introduction

Signaling pathways enable organisms to respond to environmental threats and avoid disease. The organismal phenotypes that result from the action or dysfunction of these pathways is determined by gene-environment interactions. An example of this is host-pathogen interactions, where organisms face many microbes in their environment, requiring the host immune system to defend against infection. Immunity includes both antibody-based adaptive immunity and innate immunity, which is the first line of defense. Most research on innate immunity has focused on mechanisms that reduce pathogen load, such as the regulation of anti-microbial peptides (AMPs) (Kim and Ewbank, 2018). However, less is known about the role of host metabolism in supporting survival independently of anti-bacterial responses.

*Caenorhabditis elegans*, a small free-living nematode, has been used as a model organism for decades due to its short lifespan, easy laboratory maintenance, genetic tractability and physical features (Brenner, 1974; Consortium*, 1998). The *C. elegans* diet consists of available bacteria in their environment, which in the laboratory is a non-pathogenic strain of *Escherichia coli*. This system is easily modified to study immunity, since the food source can be replaced with pathogenic bacteria, making it an excellent model system for the study of immunity (Nicholas and Hodgkin, 2002). In the wild, this soil-dwelling nematode encounters numerous pathogens, relying on its immune system for survival. Although *C. elegans* lack adaptive immunity, they have several other mechanisms for immune defense. These include pathogen avoidance behaviors and innate immunity including physical barriers and antimicrobial peptide expression. Another potential mechanism of defense is immune tolerance, which aims to reduce how an organism withstands the negative side-effects of an infection, rather than reducing the infection directly, but whether this mechanism contributes to *C. elegans* survival on pathogens has yet to be determined.

Immunity, like other aspects of metazoan physiology and development, is dependent on cell signaling. In particular, Transforming Growth Factor-β (TGF-β) signaling in multicellular animals is widely conserved and has been shown to regulate many aspects of cell function (Robertis, 2008). Bone Morphogenetic Proteins (BMP) are a major group within the TGF-β superfamily, first identified for regulating bone and cartilage development (Urist and Strates, 1971), but emerging as modulators of homeostasis. The BMP-like DBL-1 signaling pathway in *C. elegans* regulates innate immunity, lipid metabolism, body size and male tail development, among other functions (Gumienny and Savage-Dunn, 2013; Yamamoto and Savage-Dunn, 2023). This signaling pathway begins with the ligand DBL-1 (Morita et al., 1999; Suzuki et al., 1999), which binds a heterotetrameric receptor complex composed of the type I receptor SMA-6 (Krishna et al., 1999) and the type II receptor DAF-4 (Estevez et al., 1993). The signal is then transduced by the receptor-regulated Smads SMA-2 and SMA-3, and the common mediator Smad SMA-4 (Savage et al., 1996). This pathway was first identified as a major regulator of innate immunity when mutants of the DBL-1/BMP pathway exhibited decreased survival when exposed to the pathogenic bacteria *Serratia marcescens* (Mallo et al., 2002). Defects in immunity have been seen on many other pathogens, including bacteria *E. coli*, *Enterococcus fecalis*, *Pseudomonas aeruginosa* strain PA14, *Salmonella enterica*, *Salmonella typhimurium* strain SL1344*, Photorhabdus luminescens*, and the nematophagous fungus *Drechmeria coniospora* (Ciccarelli et al., 2024a; Portal-Celhay et al., 2012; So et al., 2011; Tenor and Aballay, 2008; Zugasti and Ewbank, 2009). The DBL-1/BMP signaling pathway also regulates lipid metabolism in *C. elegans*, as BMP mutants show reduced fat stores (Clark et al., 2018; Clark et al., 2021; Yu et al., 2017).

The DBL-1 regulation of immune response and lipid metabolism have, thus far, been seen as separate. However, here we explore whether these two activities are connected, and whether DBL-1 regulation of lipid metabolism has implications in immune response. We find that pathogen exposure affects fat storage, expression of genes involved in fatty acid desaturation and β-oxidation, and lipid droplet dynamics, in a DBL-1/BMP-dependent manner. Thus, BMP signaling regulates fatty acid metabolism after bacterial pathogen exposure for improved survival.

## Materials & Methods

### *C. elegans* Strains and Growth Conditions

*C. elegans* strains were grown on EZ worm plates containing streptomycin (550 mg Tris-Cl, 240 mg Tris-OH, 3.1 g Bactopeptone, 8 mg cholesterol, 2.0 g NaCl, 200 mg streptomycin sulfate, and 20 g agar per liter) to be consistent with previous studies from the lab (Ciccarelli et al., 2024a; Clark et al., 2018). All strains were maintained on *E. coli* (DA837) at 20°C. N2 was used as a wildtype control in all experiments. BMP mutant strains used were: LT207 *sma-3(wk30)*, LT121 *dbl-1(wk70)*. Lipid metabolism mutant strains used were: BX156 *fat-6(tm331); fat-7(wa36)*, BX17 *fat-4(wa14)*, BX24 *fat-1(wa9)*, BX52 *fat-4(wa14) fat-1(wa9)*, VS18 *maoc-1(hj13)*, DR476 *daf-22(m130)*, VS8 *dhs-28(hj8)*. All mutations used are strong loss-of-function or null alleles. Fluorescent lipid droplet reporter strain was LIU1 *IdrIs1 [dhs-3p::dhs-3::GFP + unc-76(+)]*, CS772 *dbl-1(wk70); IdrIs1 [dhs-3p::dhs-3::GFP + unc-76(+)]*. Genetic information was obtained from Wormbase (Sternberg et al., 2024).

### Bacteria

Control bacteria used in all experiments was *Escherichia coli* strain DA837, cultured at 37°C. Two pathogens were used for pathogenic bacteria exposure: *Serratia marcescens* strain Db11 (ATCC #13880) cultured at 37°C and *Photorhabdus luminescens* (ATCC #29999) cultured at 30°C. All experiments involving pathogens were conducted on EZ worm plates without streptomycin.

### Oil Red O (ORO) Neutral Lipid Staining

Oil Red O (ORO) staining was done as previously described (Clark et al., 2018). ORO stock solution was prepared by dissolving 0.25 g ORO powder in 50 mL isopropanol. Animals were collected after the desired time of pathogen exposure in PCR tube caps and washed three times in PBS to remove excess bacteria. Worms were fixed for 1 hour in 60% isopropanol while rocking at room temperature with caps covered with PCR tubes. While worms were fixing, the ORO working solution was made and allowed to rock at room temperature for 1 hour. After the working solution had equilibrated for 1 hour, it was filtered using a 10 mL syringe through a 0.45 µm filter, then through a 0.2 filter. The 60% isopropanol was removed and replaced with ORO working solution. The caps were covered with tubes and left overnight to stain while rocking at room temperature. The next day, the ORO was removed, and worms were washed once with PBS with 0.01% Triton, and then left in PBS while preparing slides for imaging. Worms were mounted on 2% agar pads on glass slides and imaged on a Zeiss Axioscope 2 using a Gryphax camera with Gryphax software. Images were taken using a 40x objective. Average intensity in the anterior intestine was quantified using ImageJ software.

### Survival Analysis

Survival analysis was done as previously described (Ciccarelli et al., 2024a; Ciccarelli et al., 2024b) Each survival plate was seeded with 500 µL of Db11. 50 µM 5-Fluoro-2’-deoxyuridine (FUdR) was added to each plate to prevent progeny and reduce the incidence of matricide by internal hatching of embryos. All survival experiments were carried out at 20°C. For each genotype, 120 L4 animals were picked for the experiment, and 20 animals plated per survival plate. The numbers of alive and dead animals were counted at least 4 days per week. During the experiment, some animals were lost due to burrowing, desiccation, etc. These animals were censored as their deaths were not observed. All survivals were repeated. Statistical analysis was done using Log-rank (Mantel-Cox) test.

### Bacterial Supernatant Preparation

Overnight bacterial culture of DA837 and Db11 was prepared according to temperatures specified above. Cultures were centrifuged for 5 minutes at 4000 x *g*, after which the supernatant should be relatively clear, and the bacteria should be separated into a pellet. The supernatant was filtered through a 0.45 µm pore syringe filter to remove any remaining bacterial cells in the supernatant, however, secreted peptides should be able to pass through the filter. The supernatant was added to plates already seeded with DA837 in a 1:1 volume ratio, and allowed to dry.

### RNA-Seq

Worms were synchronized with an overnight egg-lay and 4 hour timed hatch. Animals were grown on DA837 until L4, at which point they were washed with M9 buffer and transferred either to Db11 plates, or new DA837 plates. After 24 hours, they were washed with M9 again and collected in 15 mL tubes. They were washed 3x with M9, removing supernatant each time. RNA was extracted using a Trizol and chloroform precipitation, followed by Qiagen RNeasy mini kit. RNA concentration was measured using a Qubit with the RNA Broad Range kit. Samples were frozen at -80C until ready to send for sequencing. Three biologically independent replicates were collected. Sequencing was done at Azenta, resulting in a range of 22-40 M single-end reads per sample, with phred scores of 38-39. Reads were mapped to the *C. elegans* genome (WS273). Gene counts were generated with STAR. Approximately 92% of reads aligned. EdgeR was used to determine differentially expressed genes (FDR < 0.05). Analysis was done using pandas, and Venn diagrams were generated with matplotlib.

### Fluorescence Microscopy and Image Analysis

Animals were mounted on 2% agarose pads containing a 3 µL drop of 2.5 mM levamisole for immobilization. Images were taken on Zeiss LSM 900 with Airyscan 2 with Zen System software and a 63x objective. The anterior and posterior regions of the intestine were imaged as Z-stacks. In Fiji, the Z-stacks were converted to a maximum intensity projection, and the diameter and count of all visible lipid droplets in a 400 um^2^ area were measured. For each experiment, n=10 per condition was repeated in duplicate.

### Fatty Acid Supplementation

Supplementation plates were prepared by making a base of EZ worm plates with no antibiotics and adding 0.1% tergitol (NP40). They were autoclaved as usual, then after the media cooled to 50°C, 0.8 mM of fatty acid stock solution was added and allowed to stir until homogenous. Plates were then poured as usual.

### ATP/PRO

Worms were synchronized by bleaching. Animals were grown on DA837 until L4, at which point they were washed with M9 buffer and transferred either to Db11 plates, or new DA837 plates. After 24 hours, they were washed with M9 again and collected in 15 mL tubes. They were washed 3x with M9, removing supernatant each time. For each sample, 20 µL of worm pellet was transferred into a labeled eppendorf tube, 180 µL of boiled Tris-EDTA buffer added and the tube incubated at 100°C for 2 minutes. Tubes were then sonicated on ice by pulsing for 4 minutes at 60%, then centrifuged at 14,440 x *g* for 10 minutes. The supernatant was transferred to new tubes, and used for measuring ATP and protein. ATP measurement was done using the Roche ATP Bioluminescence Assay Kit. Protein measurement was done using the Thermo Scientific Pierce BCA Protein Assay Kit. Each assay was done in 96-well plates and measured with the Tecan Spark plate reader.

### Statistical Analyses

Statistical analysis was performed in Graphpad Prism 10.

## Results

### Pathogen exposure causes BMP-dependent alterations in fat storage

The first step in determining whether the immune response and lipid metabolism are connected is to identify whether there is a change in fat accumulation after pathogen exposure. While we know mutants of the DBL-1/BMP signaling pathway have low fat accumulation (Clark et al., 2018), whether the levels change after infection was unknown. *dbl-1* mutants and wildtype controls at the fourth larval stage (L4) were placed on either non-pathogenic control *E. coli* bacteria, or onto pathogenic bacteria, and after 24 hours, when animals become young adults, we conducted Oil Red O fat staining (ORO) to quantify their fat stores (Figure 1A). We selected ORO because this stain accurately corresponds to triglyceride levels measured in biochemical studies (O’Rourke et al., 2009). The 24 hour timepoint was selected as this is sufficient time for intact bacteria to proliferate in the intestinal lumen, and for some antimicrobial peptides to increase in expression (Mallo et al., 2002). We compared each condition to its internal control. In wildtype animals, we found that 24 hour exposure to a pathogen of moderate virulence, *Serratia marcescens*, resulted in a non-significant change in fat accumulation (Figure 1B). However, 24 hour exposure to a pathogen of severe virulence, *Photorhabdus luminescens*, resulted in a significant decrease in fat accumulation, with an average magnitude change of 10% (Figure 1B). In contrast to wildtype animals, *dbl-1* mutants had a dramatic and highly significant loss in total fat stores when exposed to *S. marcescens* and *P. luminescens*, with magnitudes of 25-50% (Figure 1C). Images of the *dbl-1* mutant stained animals show a substantial decrease in fat stores (Figure 1E, 1F). To validate these results in another BMP signaling mutant, we repeated the experiment with *sma-3*/*Smad* mutant animals. Similar to *dbl-1* mutants, *sma-3* mutants also had a substantial decrease in fat stores, when exposed to either *S. marcescens* or *P. luminescens* (Figure 1D). We conclude that the BMP signaling pathway regulates lipid stores during infection with a pathogen. For all further experiments, we used *S. marcescens* as a pathogen because the difference between wildtype and BMP mutants was most significantly different on the moderate-virulence pathogen.

**Figure 1.**
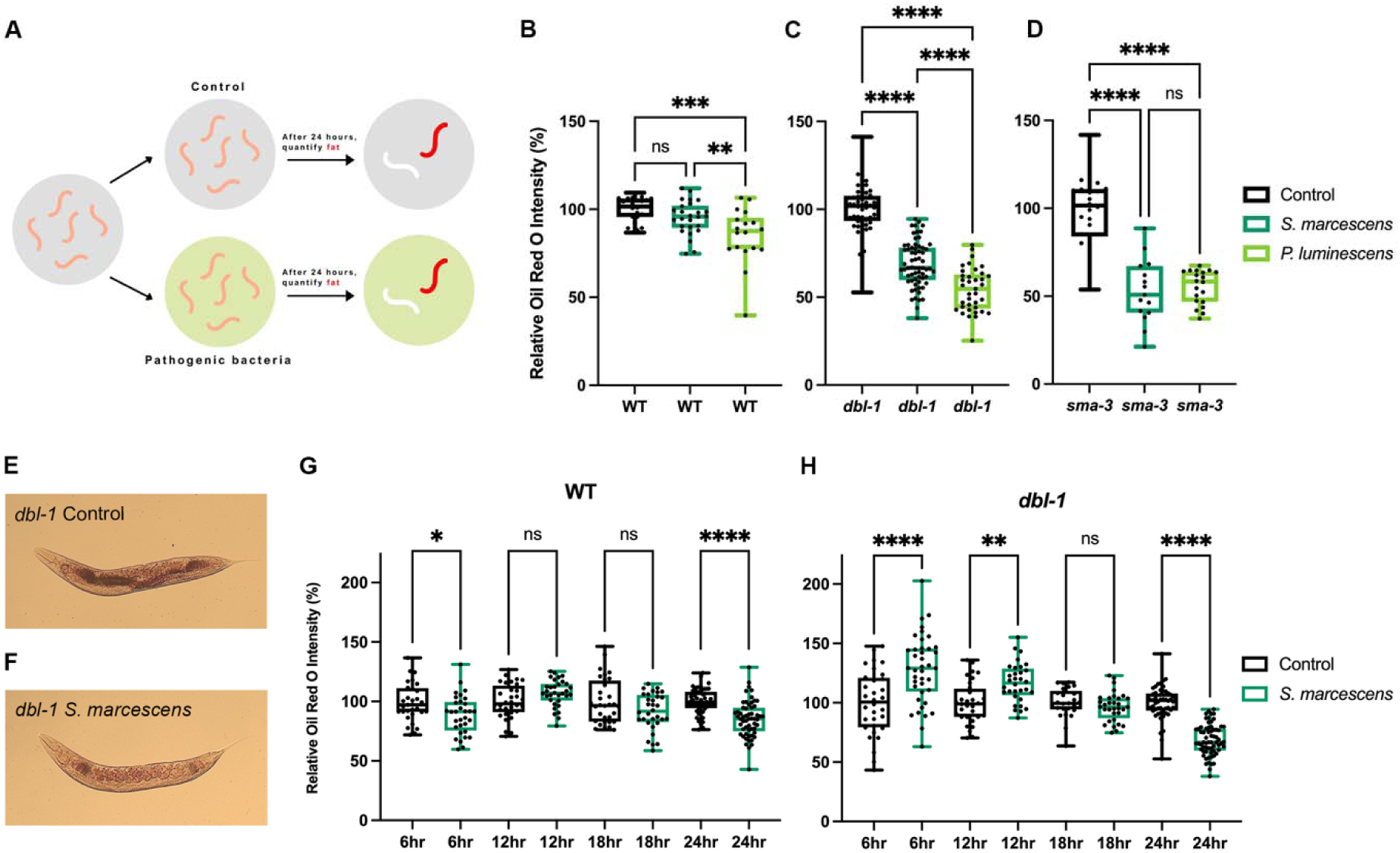
Pathogen exposure causes BMP-dependent alterations in fat storage. (A) Experimental schematic: L4 animals are transferred to either control or pathogenic bacteria. After 24 hours, the animals are stained with Oil Red O (ORO). (B, C, D) Lipid accumulation of wildtype (WT), *dbl-1* and *sma-3* animals, respectively, after 24 hour pathogen exposure, stained with ORO. ORO experiments were repeated in triplicate on independent biological samples, with at least 15 animals per condition. Brown-Forsythe and Welch ANOVA multiple comparisons tests were used to determine significance. (E, F) Representative images of lipid accumulation in *dbl-1* mutants after 24 hour exposure to *E. coli* or *S. marcescens*, respectively. (G, H) Lipid accumulation of WT and *dbl-1*, respectively, after exposure to *E. coli* or *S. marcescens* for 6 hours, 12 hours, 18 hours, or 24 hours. ORO experiments were repeated in triplicate on independent biological samples, with at least 30 animals per condition. Brown-Forsythe and Welch ANOVA multiple comparisons tests were used to determine significance. ns = p > 0.01; * = p<=0.05; ** = p<=0.01; *** = p<=0.001; **** = p<=0.0001.

We were surprised to see the dramatic fat loss in the BMP mutants in only 24 hours of pathogen exposure. We wondered whether 24 hours was the earliest timepoint at which the decrease would be observed, and thus, we investigated the temporal dynamics of lipid accumulation upon infection prior to 24 hours. We repeated the ORO fat staining after pathogen exposure, and assayed lipid levels at 6 hours, 12 hours, 18 hours, and 24 hours in wildtype and *dbl-1* mutants. In wildtype, across all time points, there was either no significant change in fat after pathogen exposure, or there was a small decrease, indicating that wildtype animals are able to maintain lipid homeostasis (Figure 1G). However, in *dbl-1* mutants, we see a more dynamic trend unfold (Figure 1H). At 6 hours, animals exposed to pathogen showed a highly significant increase in fat stores. At 12 hours, there was still an increase in fat stores, however the magnitude was smaller. At 18 hours, it appeared there was no difference between animals on control bacteria and the pathogenic *S. marcescens*. At 24 hours, we observed the significant decrease we had originally observed. We conclude that in *dbl-1* mutants, there is an immediate increase in fat stores in response to pathogen exposure, which are depleted by 24 hours. These results suggest upon pathogen exposure, the loss of BMP signaling results in lipid dysregulation, perhaps having consequences for these animals’ survival.

### Alterations in fat storage do not require pathogen ingestion

Given that *C. elegans* are bacteriotrophs, and consume bacteria in their environment as food, we wondered whether the dramatic changes in fat exhibited by BMP mutants after pathogen exposure was simply due to nutritional differences between *E. coli* and bacterial pathogens. To test this, we determined the effect of exposure to the supernatant of the bacterial culture, made by centrifuging and filtering the overnight bacterial culture. We expect that this would remove all bacterial cells, and thus the majority of the nutrition, while leaving supernatant with secreted peptides, a potential source of pathogenicity. We first conducted a survival assay, to confirm that the filtered supernatant had pathogenic effects. All the plates had a lawn of *E. coli*, as a food source for the animals, supplemented with either the *E. coli* or *S. marcescens* filtered supernatant on top. We found that both wildtype and *dbl-1* mutant animals experienced a shorter lifespan when placed on the *S. marcescens* supernatant compared with the *E. coli* supernatant (Figure 2A). This suggests that the supernatant retains some pathogenicity, though it is milder than live bacteria. The decreased survival was more dramatic in *dbl-1* mutants, as expected, given that they are known to have an impaired immune response (Figure 2B) (Mallo et al., 2002). Since the *S. marcescens* filtered supernatant retained pathogenicity, but removed nutritional changes, we repeated the 24 hour exposure followed by ORO fat staining. Both wildtype and *dbl-1* mutants had changes in fat accumulation following the filtered supernatant exposure, consistent with it triggering an organismal response. Wildtype animals showed a decrease in fat levels when exposed to the *S. marcescens* supernatant, more so than when a regular lawn of *S. marcescens* was employed (Figure 2C). In *dbl-1* mutants, we observed an increase in fat stores at 24 hours, similar to that observed after 6-12 hours of exposure to a lawn of *S. marcescens* (Figure 2D). Taken together, these results confirmed that the changes in fat accumulation cannot solely be due to nutritional changes.

**Figure 2.**
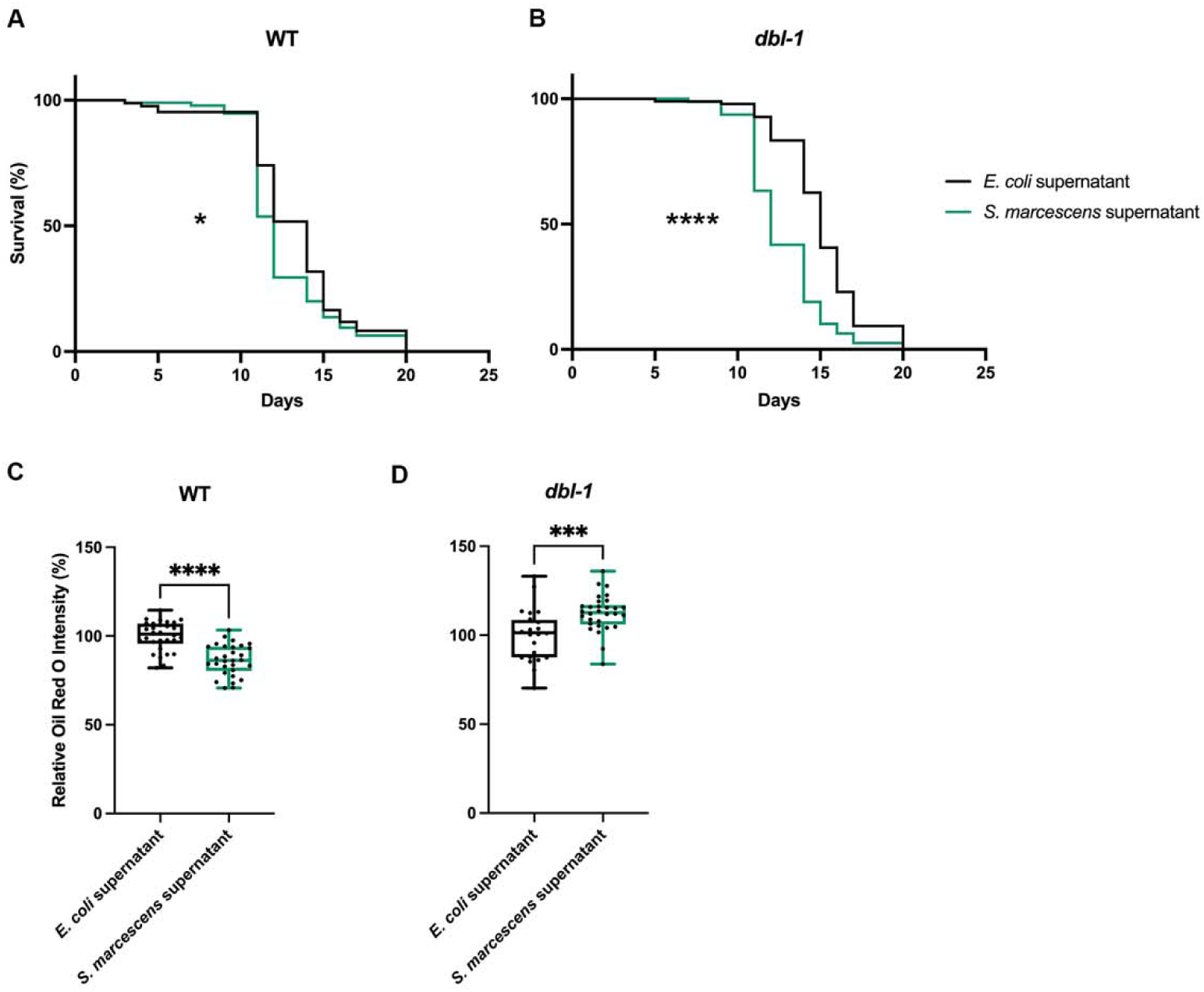
Alterations in fat storage are due to pathogenicity. (A) Survival analysis of wildtype animals on *E. coli* bacteria, with either *E. coli* supernatant or *S. marcescens* supernatant. N values: *E. coli* supernatant (85), *S. marcescens* supernatant (95). (B) Survival analysis of *dbl-1* animals on *E. coli* bacteria, with either *E. coli* supernatant or *S. marcescens* supernatant. N values: *E. coli* supernatant (96), *S. marcescens* supernatant (79). (C) Lipid accumulation of wildtype animals after 24 hour exposure to *E. coli* bacteria, with either *E. coli* supernatant or *S. marcescens* supernatant, stained by ORO. ORO experiment was repeated in duplicate on independent biological samples, with 30 animals per condition. Brown-Forsythe and Welch ANOVA multiple comparisons tests were used to determine significance. (D) Lipid accumulation of *dbl-1* mutant animals after 24 hour exposure to *E. coli* bacteria, with either *E. coli* supernatant or *S. marcescens* supernatant, stained by ORO. ORO experiment was repeated in duplicate on independent biological samples, with 30 animals per condition. Brown-Forsythe and Welch ANOVA multiple comparisons tests were used to determine significance. ns = p > 0.01; * = p<=0.05; ** = p<=0.01; *** = p<=0.001; **** = p<=0.0001.

### RNA-seq reveals that lipid metabolism genes are highly overrepresented among differentially expressed genes in response to *S. marcescens*

Our results suggested an active organismal response to pathogen exposure leading to changes in lipid stores, thus we hypothesized that transcriptional changes could be responsible for the changes in lipid accumulation observed after pathogen exposure. We conducted whole animal RNA-seq of wildtype and *dbl-1* mutants after 24 hour exposure to either *E. coli* or *S. marcescens*. We then compared the transcriptional response on pathogen between the two genotypes, as we aimed to identify BMP-dependent genes that were differentially expressed under pathogenic conditions. Differentially expressed genes (DEGs) were identified as those up-regulated or down-regulated in response to pathogen (FDR < 0.05). We did not see a strong pattern of activation or repression in either genotype (Figure 3A, 3B). Some DEGs were shared in both genotypes, which are likely part of a conserved response that is not BMP-dependent. The largest category of DEGs were those upregulated in WT (122 genes), but not in *dbl-1* mutants (Figure 3C). This category represents pathogen response genes that are BMP-dependent.

**Figure 3.**
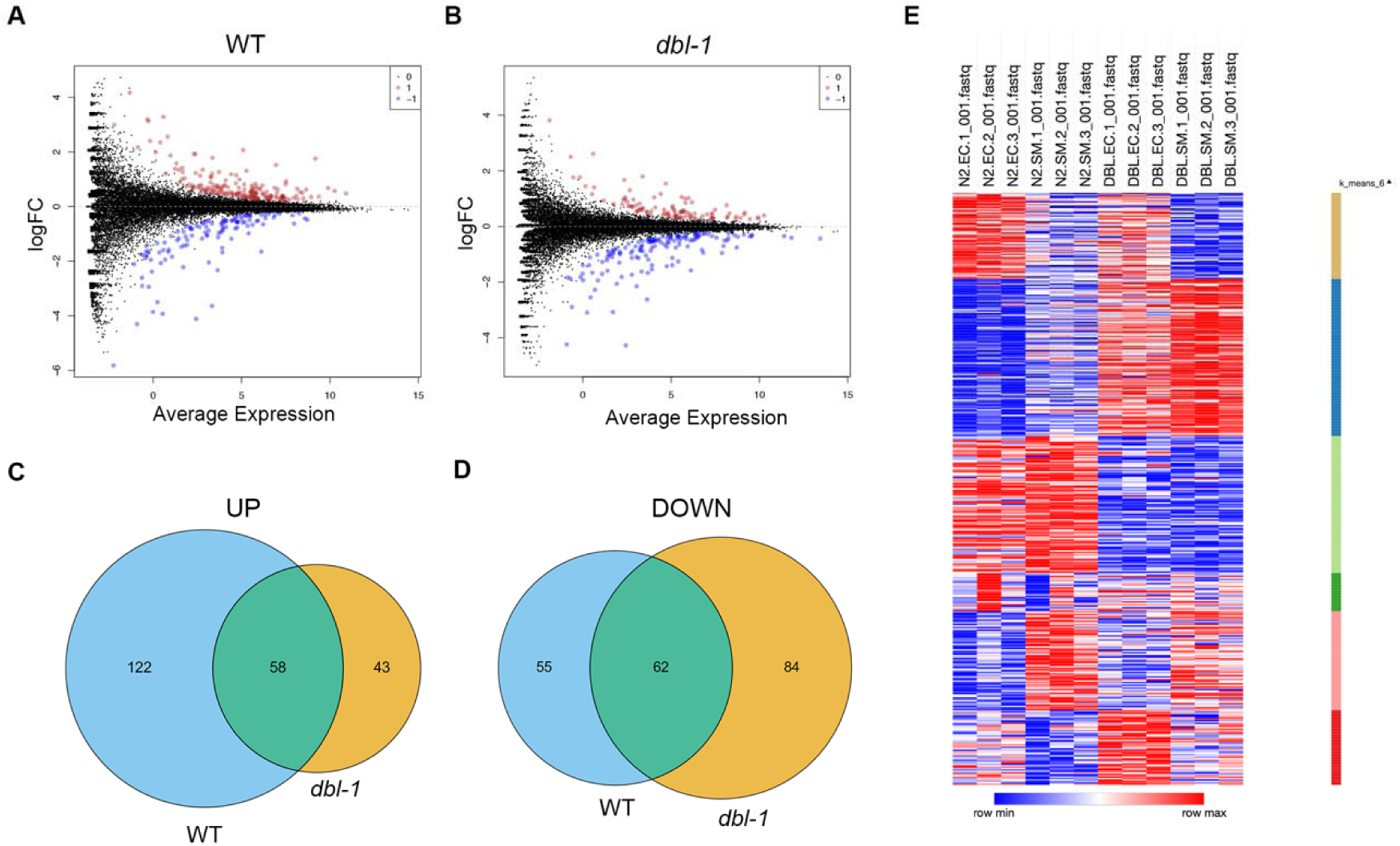
Transcriptional changes in WT and *dbl-1* mutants after pathogen exposure. (A, B) Volcano plots of RNA-seq log fold change versus average expression for individual genes on pathogen compared to control bacteria, in wildtype animals and *dbl-1* mutants, respectively. Data points labeled in red are upregulated DEGs, and data points labeled in blue are downregulated DEGs. (C, D) Venn diagrams of upregulated and downregulated DEGs, respectively, between wildtype and *dbl-1* mutants. (E) Heatmap of RNA-seq results, with each row being the relative expression of an individual gene. Genes with similar patterns of expression are grouped by K-means clustering. Figure generated using Morpheus by Broad Institute (RRID: SCR_017386).

We investigated the DEGs further by employing WormCat (Holdorf et al., 2020) to identify enriched gene sets and to generate a heatmap for visualization. We anticipated that the stress response, specifically the pathogen stress response, would be highly enriched, which we observed in both genotypes. We also hypothesized that several lipid metabolism genes would appear in the DEGs, given our previous experiments. Strikingly, lipid metabolism was the most enriched category in both genotypes (Figure 4A). If we examine the 122 genes that are upregulated in WT, and thus BMP-dependent, we observe the molecular function category is highly enriched, more so than any other molecular function category across all columns (Figure 4B). This suggests that the first 24 hours of pathogen exposure elicits a greater transcriptional response of genes involved in lipid metabolism genes than that of genes involved in the immune response.

**Figure 4.**
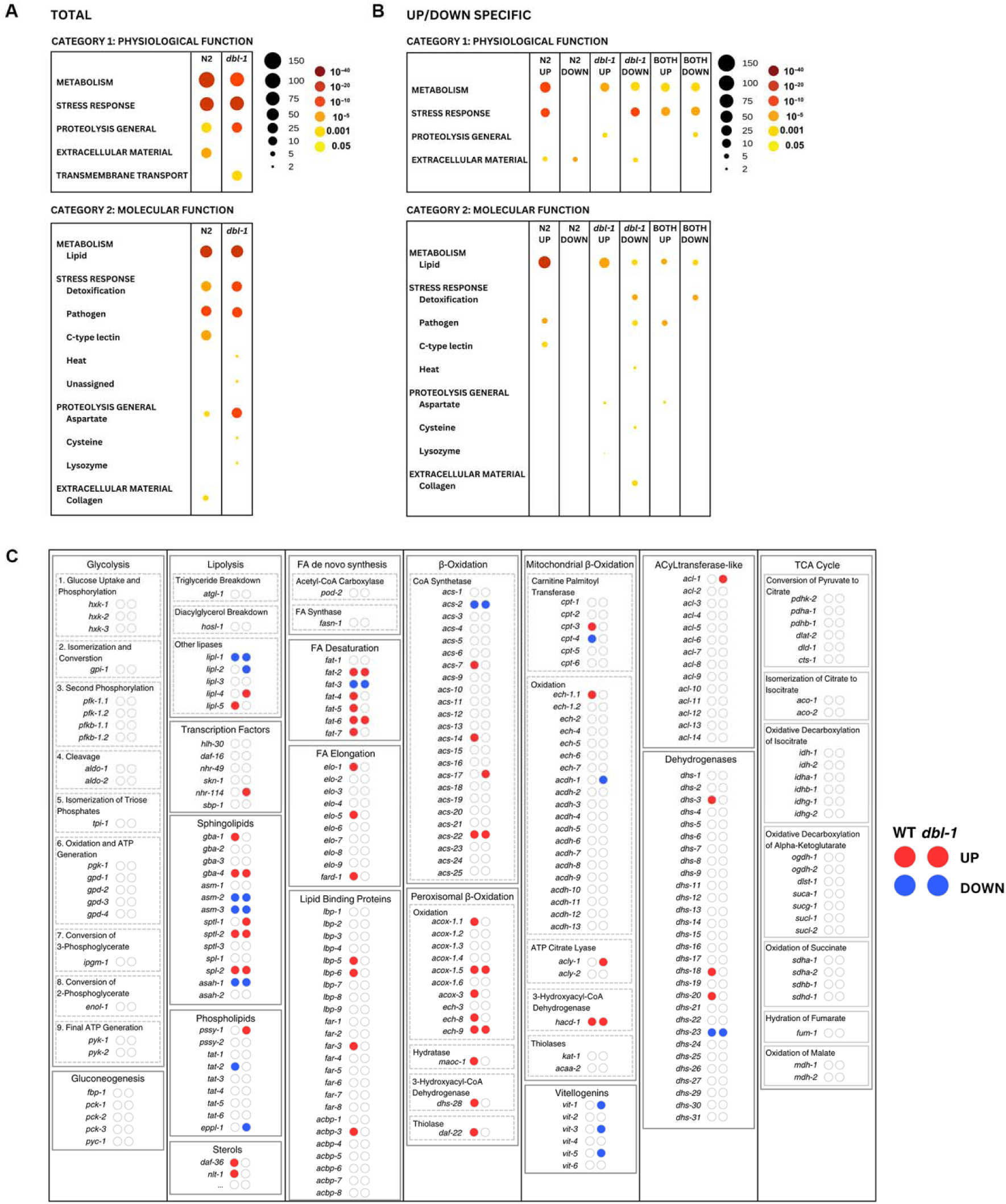
Differential gene expression for key lipid metabolism genes in *C. elegans*. (A) WormCat analysis of identified gene category enrichments, showing results per genotype. The legend shows that size of the circle represents the number of genes in the category, and the color represents P-value. (B) WormCat visualization of identified gene category enrichments, showing results for direction-specific groups of DEGs. (C) Visualization of DEGs identified by RNA-seq in the context of key genes in *C. elegans* lipid metabolism, grouped by process. There are two bubbles to the right of every gene name, indicating whether they are up- or down-regulated in wildtype animals or *dbl-1* mutants, respectively. Bubbles in red indicate upregulated DEGs and bubbles in blue indicate downregulated DEGs.

We were interested to explore which specific lipid metabolism genes were differentially expressed in wildtype and *dbl-1* mutants. First, we created a schematic to help our visualization that contained many fundamental lipid metabolism processes, with rate-limiting or important genes listed below each process. We listed the lipid metabolism genes identified in WormCat and labeled our table with whether they were differentially expressed, and if so, in which direction (Figure 4C). Some processes seem unaffected by the short exposure to pathogen, such as glycolysis and the TCA cycle. In contrast, several processes were more affected, particularly in wildtype, such as fatty acid desaturation and elongation, and β-oxidation. We hypothesize that in the presence of BMP signaling, there is an increase in these processes in response to pathogen exposure.

### Lipid droplets experience flux after pathogen exposure

We wanted to test whether changes to these processes were evident on a cellular level. Lipid droplets are sensitive to changes in both fatty acid desaturation and β-oxidation. In fatty acid desaturase mutants, lipid droplets are smaller and decreased in number, due to impaired storage (Brock et al., 2007). In β-oxidation mutants, lipid droplets are larger due to a disruption in fatty acid breakdown (Zhang et al., 2010). We selected the lipid droplet reporter DHS-3::GFP and crossed in the lipid droplet reporter to *dbl-1* loss-of-function mutants. We compared DHS-3-positive lipid droplets in wildtype and *dbl-1* animals after 24 hours of bacteria exposure. We found that lipid droplets in the anterior intestine of wildtype animals on *S. marcescens* displayed a significant increase in the number of lipid droplets, however no significant change in droplet diameter (Figure 5A, 5C). We saw a flipped trend in the posterior intestine of wildtype animals, which displayed no significant change in the number of lipid droplets, however an increase in droplet diameter (Figure 5B, 5D). *dbl-1* animals displayed no significant change in the anterior or posterior intestine, neither in lipid droplet quantity nor diameter (Figure 5A, 5B, 5C, 5D). We were surprised that pathogen exposure did not cause a reduction in lipid droplet number in *dbl-1* mutants. It is possible that expression of dehydrogenase DHS-3::GFP in these strains lowers the baseline level of lipid droplets such that further depletion in *dbl-1* mutants is not easily quantifiable.

**Figure 5.**
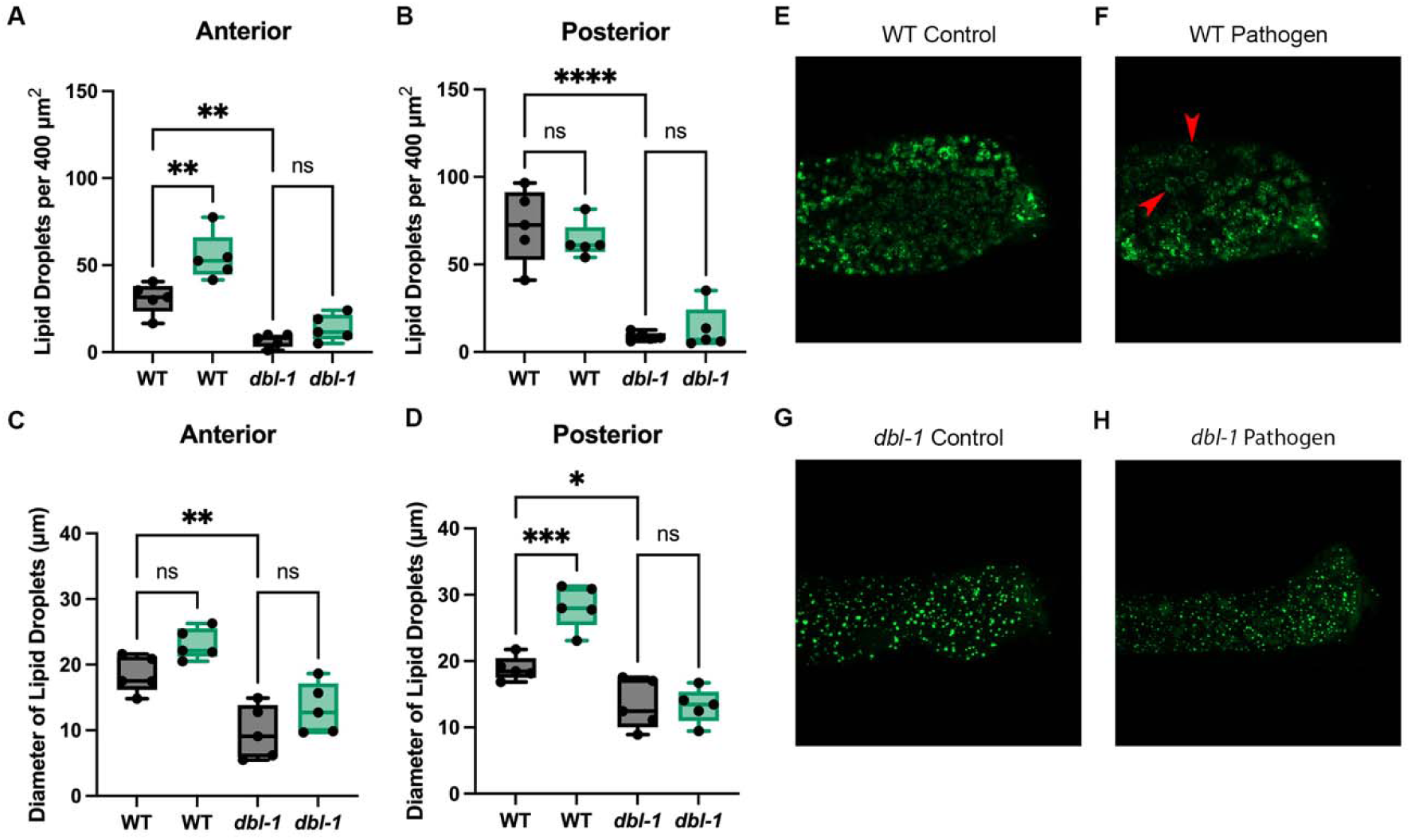
Lipid droplet dynamics are in flux after pathogen exposure. (A, B) Box-and-whisker plots of the number of lipid droplets per 400 µm^2^ in the anterior and posterior intestine. (C, D) Box-and-whisker plots of lipid droplet diameter (µm) in the anterior and posterior intestine. Brown-Forsythe and Welch ANOVA multiple comparisons tests were used to determine significance. (E, F) Representative confocal images of DHS-3::GFP after 24 hours of control bacteria exposure, and *S. marcescens* exposure, respectively. Red arrowheads point to enlarged lipid droplets. (G, H) Representative confocal images of *dbl-1*; DHS-3::GFP after 24 hours of control bacteria exposure, and *S. marcescens* exposure, respectively. Confocal experiment was repeated in duplicate on independent biological samples, with 5 animals per condition. ns = p > 0.01; * = p<=0.05; ** = p<=0.01; *** = p<=0.001; **** = p<=0.0001.

### β-oxidation plays a limited role in pathogen survival

We have found that pathogen exposure results in a lipid metabolism response, which upregulates both fatty acid desaturation and β-oxidation, and causes significant changes in lipid droplet dynamics. We next wanted to determine whether these changes impact survival on pathogenic bacteria. We first focused on β-oxidation, which we reasoned could be upregulated to convert lipid stores to the energy necessary for the immune response, as previously suggested (Dasgupta et al., 2020). We tested whether the impairment of key β-oxidation genes impacted *C. elegans* survival. We investigated three key β-oxidation genes: *maoc-1*, *daf-22* and *dhs-28*. Mutations in any of these three genes result in an increase in lipid droplet size (Li et al., 2016). *maoc-1* mutant animals showed minimal defects in survival during a *S. marcescens* survival assay (Figure 6A). *daf-22* and *dhs-28* mutants, on the other hand, had significant defects in survival (Figure 6B, 6C). *maoc-1* encodes the enzyme at the upstream rate-limiting step, while DAF-22 and DHS-28 function downstream of MAOC-1 in β-oxidation. DAF-22 and DHS-28 also have their pleiotropic roles; for example, DAF-22 and DHS-28 are both involved in the synthesis of the dauer pheromone (Butcher et al., 2009). It is possible that the involvement of DAF-22 and DHS-28 in a synthetic pathway may be responsible for the survival defects. If β-oxidation is needed for energy generation in response to the infection, then we would expect that ATP stores would be lowered upon infection, similarly to the effect we see on fat stores. We quantified the amount of ATP in each experimental condition, after 24 hours of pathogen exposure. Due to the body size phenotype of *dbl-1* mutants, we normalized each ATP concentration to the concentration of protein (PRO) in that sample. We found that there was no significant change in ATP/PRO in either genotype (Figure 6D). Thus, energy stores are maintained to a similar degree in wildtype and mutant animals, despite the significant differences in lipid metabolism between these two genotypes.

**Figure 6.**
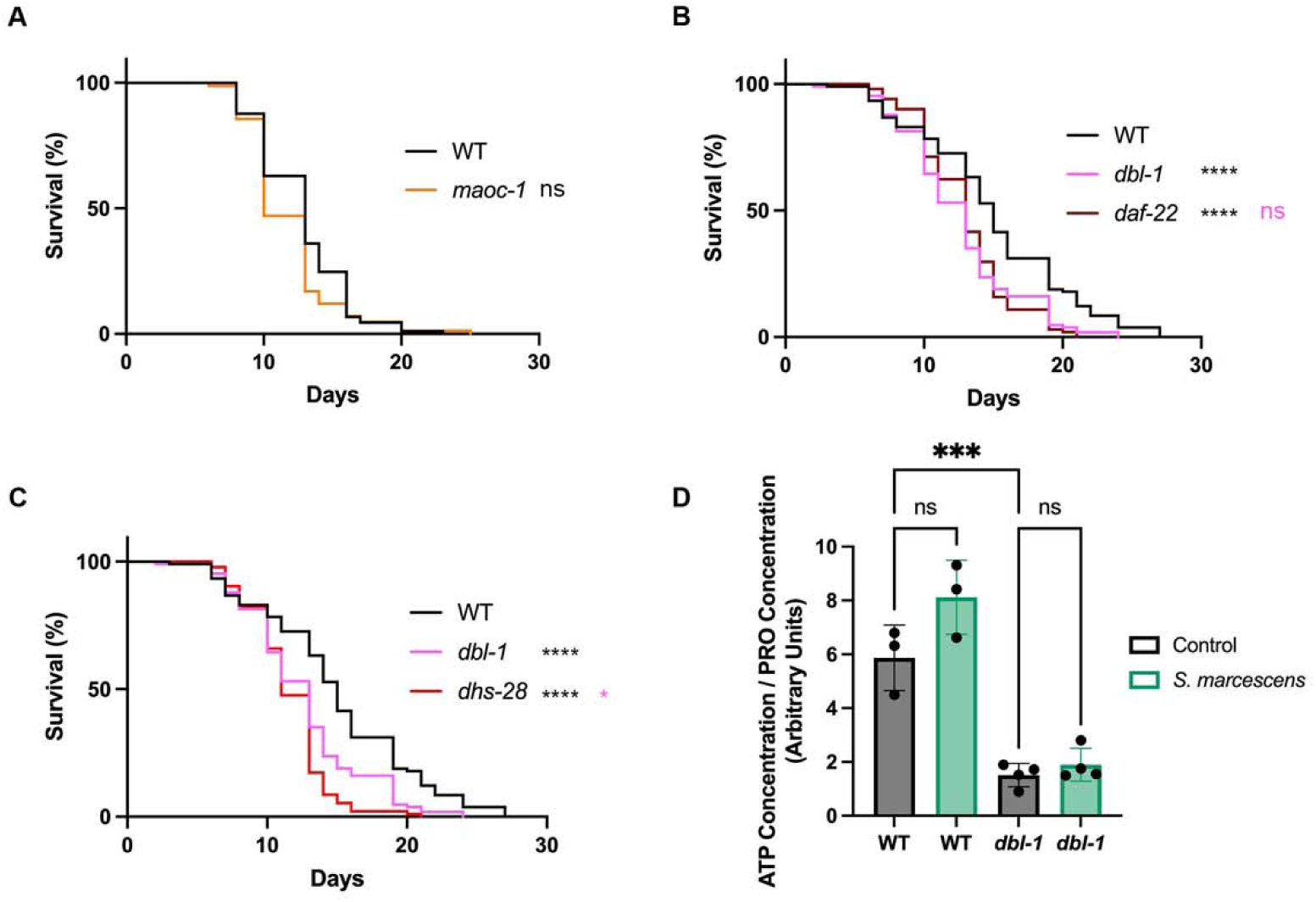
Fatty acid β-oxidation plays a minor role in survival after pathogen exposure. (A) Survival analysis of wildtype animals and *maoc-1* mutants on *S. marcescens* bacteria. N values: WT (89), *maoc-1* (83). (B) Survival analysis of wildtype, *dbl-1* and *daf-22* animals on *S. marcescens* bacteria. N values: WT (106), *dbl-1* (106), *daf-22* (101). (C) Survival analysis of wildtype, *dbl-1* and *dhs-28* animals on *S. marcescens* bacteria. N values: WT (106), *dbl-1* (106), *dhs-28* (93). (D) Ratio of ATP concentration to protein concentration in wildtype and *dbl-1* mutants after 24 hour exposure to either control *E. coli* or pathogenic *S. marcescens*. ns = p > 0.01; * = p<=0.05; ** = p<=0.01; *** = p<=0.001; **** = p<=0.0001. Black asterisks denote significance relative to wildtype control; pink is significance relative to *dbl-1*.

### Fatty acid desaturases promote survival in response to pathogen

Our results demonstrated that β-oxidation does not play a major role in the survival of animals after pathogen exposure, thus we next tested the role of genes involved in lipid synthesis. We focused on genes that function in fatty acid desaturation, hypothesizing that these genes are upregulated in WT to increase lipid synthesis under pathogenic conditions. FAT-6 and FAT-7 encode redundant Δ9 desaturases and are responsible for converting stearic acid into oleic acid, a rate-limiting step in lipid synthesis (Brock et al., 2006). *fat-6;fat-7* mutants show a severe decrease in survival when exposed to pathogen, very similar to the level observed in *dbl-1* mutants (Figure 7A). These mutants are defective in converting stearic acid to oleic acid, thus we hypothesized that supplementing animals with oleic acid would rescue the survival defect. In WT, the addition of 0.8 mM oleic acid had no effect on survival (Figure 7B). However, *dbl-1* mutants displayed improved survival, when exposed to pathogen and the addition of 0.8 mM oleic acid improved survival, consistent with our hypothesis (Figure 7C).

**Figure 7.**
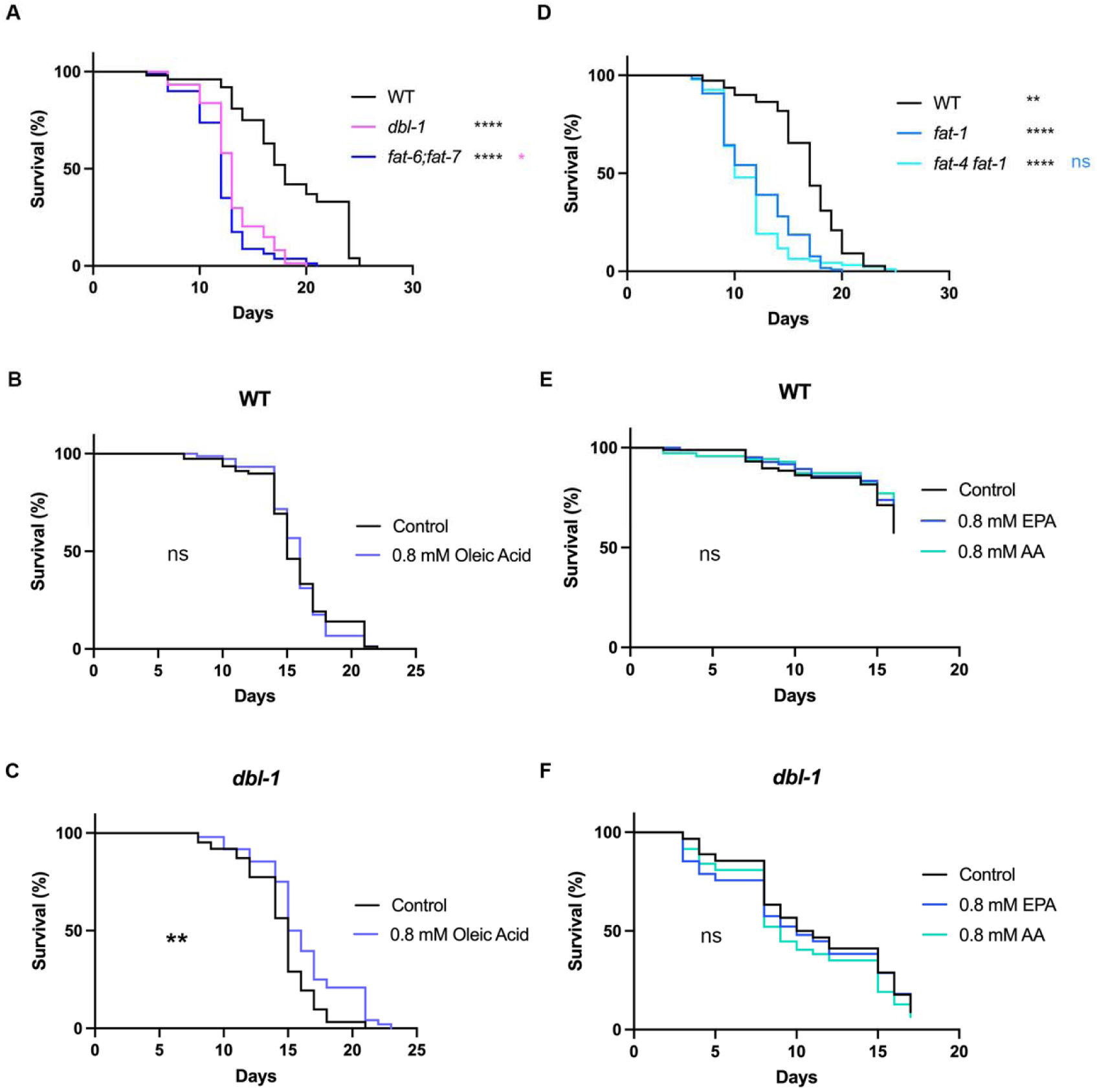
Fatty acid desaturation plays a major role in survival after pathogen exposure. (A) Survival analysis of wildtype, *dbl-1* and *fat-6;fat-7* animals on *S. marcescens* bacteria. N values: WT (100), *dbl-1* (74), *fat-6;fat-7* (80). (B, C) Survival analysis of wildtype and *dbl-1* animals, respectively, on *S. marcescens* bacteria with and without supplementation of 0.8 mM oleic acid. N values: WT Control (78), WT Oleic Acid (74), *dbl-1* Control (62), *dbl-1* Oleic Acid (48). (D) Survival analysis of wildtype, *fat-1* and *fat-4 fat-1* animals on *S. marcescens* bacteria. N values: WT (110), *fat-1* (118), *fat-4 fat-1* (94). (E, F) Survival analysis of wildtype and *dbl-1* animals, respectively, on *S. marcescens* bacteria with and without supplementation of 0.8 mM eicosapentanoic acid (EPA) or 0.8 mM arachidonic acid (AA). N values: WT Control (87), WT EPA (84), WT AA (70), *dbl-1* Control (90), *dbl-1* EPA (94), *dbl-1* AA (94). ns = p > 0.01; * = p<=0.05; ** = p<=0.01; *** = p<=0.001; **** = p<=0.0001. Black asterisks denote significance relative to wildtype control; pink is significance relative to *dbl-1*; blue is significance relative to *fat-1*.

Our experiments suggest that monounsaturated fatty acids (MUFAs) can partially, but not fully rescue, the survival of BMP mutants. We considered the possibility that polyunsaturated fatty acids (PUFAs) may also be required for survival upon exposure to pathogen. In support of that, FAT-4, and several elongases, are also induced in animals exposed to pathogen and this response is BMP-dependent. We chose to look at the *fat-4 fat-1* double mutant, which would eliminate most PUFAs. We found that *fat-4 fat-1* had a significant defect in survival after pathogen exposure (Figure 7D). We concurrently assayed the survival of *fat-4 fat-1* with *fat-6;fat-7* mutants to determine whether one was more severe, and we found no significant difference between the two double mutants (Figure S1). We conclude that disruption at any point in fatty acid desaturation has a strong impact on the survival to pathogen exposure. We were curious whether supplementing with single PUFAs may also rescue the survival defect. Since FAT-4 is a Δ5 desaturase that produces the PUFAs arachidonic acid and eicosapentanoic acid (Watts and Browse, 2002), we supplemented with 0.8 mM arachidonic acid or 0.8 mM eicosapentanoic acid. Surprisingly, we observed no change to the survival of wildtype or *dbl-1* animals. Therefore, unlike MUFAs, single PUFAs are not sufficient for rescuing *dbl-1*’s impaired immune response.

## Discussion

We have found that after a short exposure to the bacterial pathogen *Serratia marcescens*, *C. elegans* undergo BMP-dependent changes in lipid metabolism flux, both in the synthesis and breakdown of lipids. We find that these changes are associated with the induction of genes encoding proteins involved in β-oxidation and fatty acid desaturation. Furthermore, pathogen exposure of wildtype animals induces an increase in lipid droplet diameter, but a decrease in lipid droplet number. In wildtype animals, the net effect of these two changes is the maintenance of total fat stores, whereas *dbl-1*/BMP loss-of-function mutants exhibit first an increase, and then a decrease in fat stores. Since both genetic backgrounds maintain a constant concentration of ATP following pathogen exposure, we believe that these changes may allow for the generation of MUFAs, rather than the generation of ATP. Consistent with our results, oleic acid was found to be required for a normal immune response, with *fat-6; fat-7* mutants deficient in oleic acid having a decreased survival when exposed to bacterial pathogens, such as *Pseudomonas aeruginosa* and *Serratia marcescens* (Anderson et al., 2019). Furthermore, another paper also observed a pathogen-specific alteration of fat stores in wildtype animals, with *P. aeruginosa, S. aureus, E. faecalis,* and *C. neoformans* depleting neutral lipids after 8 hours of exposure, while similar exposure to *S. typhimurium* did not (Dasgupta et al., 2020).

An organism’s immune response employs many strategies in concert to fight illness and infection. These strategies can be clustered into three approaches: pathogen avoidance, immune resistance, and immune tolerance (Medzhitov et al., 2012). Pathogen avoidance occurs prior to the organism making contact with a pathogen, and aims to reduce the risk of exposure to infection. This typically manifests as a physical distancing of an organism from a potential pathogen. Early work in pathogen avoidance can be attributed to rodent models, and even wild populations of lobster (Behringer et al., 2006; Kavaliers et al., 2004). In humans, the emotion of disgust is central to pathogen avoidance, as this core emotion is triggered by potential pathogens, or pathogen-harboring substances (Curtis, 2014). In *C. elegans*, pathogen avoidance manifests as animals physically distancing themselves from pathogenic bacteria in their petri dish environment, often climbing up the plastic sides and desiccating, or burrowing into the agar. *dbl-1* mutants have an increased avoidance of *E. coli*, suggesting that the standard lab food source for *C. elegans* may have increased pathogenicity in these animals compared to wildtype animals (Madhu et al., 2023; Olofsson, 2014). *dbl-1* mutants also show increased avoidance to three Gram-negative bacteria, compared to wildtype animals (Madhu et al., 2023), demonstrating the DBL-1 pathway is required to suppress avoidance behavior.

The next strategy is immune resistance, which encompasses most traditional notions of disease-fighting, and can be found, to some extent, in all organisms. Immune resistance aims to reduce the pathogen burden once an infection is already established. This approach includes both innate immunity, such as physical barriers (skin, etc.) and the upregulation of AMPs, as well as adaptive immunity, such as antibodies. *C. elegans* only have innate immunity, thus the primary method of resistance is the upregulation of AMPs in response to pathogen exposure. DBL-1/BMP signaling regulates the expression of many immune response genes, including lectins, lysozymes, lipases, P-glycoproteins (PGPs) of the ATP-binding cassette transporter family, caenacin AMPs, and saposin-like proteins (Alper et al., 2007; Liang et al., 2007; Madhu et al., 2020; Mallo et al., 2002; Mochii et al., 1999; Roberts et al., 2010; Zugasti and Ewbank, 2009). Among these, caenacins play a critical role in the immune response, and are induced upon infection. Notably, DBL-1 signaling promotes *cnc-2* expression in the epidermis in a dose-dependent manner (Zugasti and Ewbank, 2009). Recent studies also revealed that CNC-2, along with another AMP, ABF-2, are regulated by SMA-3 activity in the pharynx (Ciccarelli et al., 2024a). Conversely, DBL-1 signaling negatively regulates the expression of the saposin-like protein SPP-9 (Madhu et al., 2020; Roberts et al., 2010).

The last strategy in the immune response, and the least understood, is immune tolerance, which aims to reduce the negative impacts of infection on host fitness. While the presence of a pathogen in a host has direct consequences such as cell death, there are also indirect consequences that can hinder an effective immune response, such as high inflammation. Foundational work in maize and wheat (Caldwell et al., 1958; Schafer, 1971), as well as in rodent models have differentiated tolerance from resistance (Ayres and Schneider, 2008; Råberg et al., 2009; Read et al., 2008). This work allows the hypothesis to form that some genotypes may be more capable of withstanding the side effects of infection, and thus more tolerant. In *C. elegans*, processes with a role in immune tolerance may include the microbiome and lipid metabolism, however studies on these topics are less abundant than those on immune resistance. In this study, we sought to explore whether DBL-1 signaling regulates lipid metabolism under pathogenic conditions, and whether this is protective, potentially relating its effects to immune tolerance.

Based on our results, we see two possible models, which are not mutually exclusive, for the identified changes in lipid metabolism: 1) the lipid changes feed into the immune response in a way that is intended to directly combat bacteria, perhaps through some sort of systemic signaling, or 2) the lipid changes contribute to immune tolerance. To differentiate between these models, expression levels of AMPs, or bacterial load, in fat metabolism mutants could be determined.

Other work in *C. elegans* has also linked lipid metabolism and immunity. A study found that animals exposed to some pathogenic bacteria undergo lipolysis and rapidly utilize lipid droplets, regulated by nuclear hormone receptor NHR-49 (Dasgupta et al., 2020). Another study found that PUFAs, specifically gamma-linolenic acid and stearidonic acid, are required to maintain p38 MAP kinase pathway activity, and when deficient, result in an increased susceptibility to infection (Nandakumar and Tan, 2008). Another publication demonstrated that immunity-linked genes, genes upregulated in response to infection, are also often upregulated in response to lipid metabolism disruptions, both considered environmental stresses (Fanelli et al., 2023). The connecting mechanism is that these immunity-linked genes support secretory functions under stressful conditions.

In conclusion, our research has established a direct link between lipid metabolism and pathogen survival in *C. elegans,* revealing that BMP-dependent regulation of lipid stores contributes to *S. marcescens* resistance. The connection between the immune response and lipid metabolism is likely to be conserved. Our findings are consistent with work in *D. melanogaster*, where a study found that infection activates mobilization of host lipid stores, improving survival (Deshpande et al., 2022a; Deshpande et al., 2022b). Similar observations have been made in grapevines, where lipid signaling regulates pathogen response (Laureano et al., 2021), demonstrating how widespread the relationship may be across phylogenetic kingdoms.

Furthermore, in mammalian cells, there is an established requirement for lipids in fighting infection. Adipose tissue has been identified as a key contributor to the immune system by storing immune cells (Desruisseaux et al., 2007). Individual adipocytes have been implicated due to their potential regulatory role on the immune system through the secretion of hormones. Fat cells also dynamically move to wound sites and act collaboratively with macrophages to prevent infection (Franz et al., 2018). This relationship may explain the findings from human patients, as individuals that have metabolic syndromes, including 450 million diabetes patients, suffer from an increased risk for severe infection (Carey et al., 2018). Our findings thus have broad implications for understanding host-pathogen interactions and may pave the way for the development of therapies that improve outcomes against infectious diseases, particularly in the context of metabolic diseases.

## Data Availability

Files for RNA-seq are available at NIH/NCBI GEO through accession number GSE291387.

## Acknowledgements

This work was supported by NIH awards R15GM112147, R21AG075315, and R35GM153390 to CSD. Some strains were provided by the Caenorhabditis Genetics Center, which is supported by the National Institutes of Health Office of Research Infrastructure Programs (P40 OD010440). We thank Dr. Alicia Meléndez, Dr. Venkatakrishna Lappasi Mohanram, and Allen Sun for helpful comments on the manuscript. This work was carried out in partial fulfillment of the requirements for the Ph.D. degree from the Graduate Center of the City University of New York (KKY).

---

**Figure S1.**
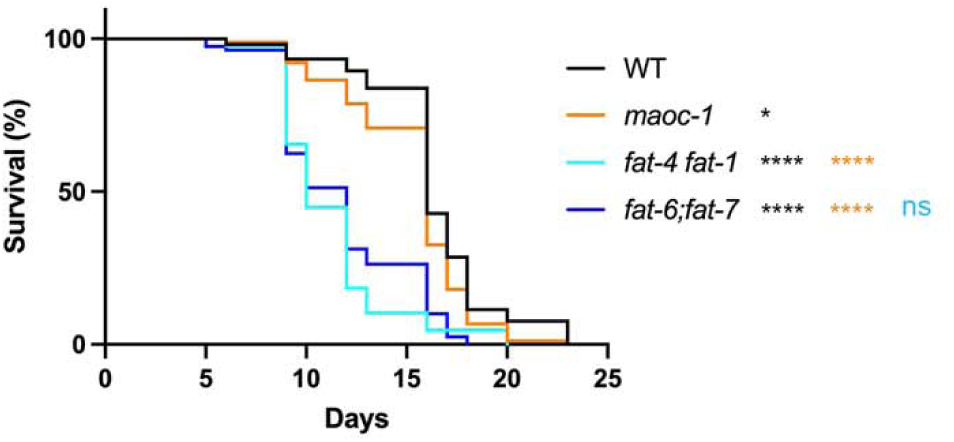
Fatty acid desaturation is more involved in survival after pathogen exposure than β-oxidation. Survival analysis of wildtype, *maoc-1*, *fat-4 fat-1* and *fat-6;fat-7* animals on *S. marcescens* bacteria. N values: WT (105), *maoc-1* (89), *fat-4 fat-1* (87), *fat-6;fat-7* (80). ns = p > 0.01; * = p<=0.05; ** = p<=0.01; *** = p<=0.001; **** = p<=0.0001. Black asterisks denote significance relative to wildtype control; orange is significance relative to *maoc-1*; teal is significance relative to *fat-4 fat-1*.

## References

Alper, S., McBride, S. J., Lackford, B., Freedman, J. H. and Schwartz, D. A. (2007). Specificity and Complexity of the Caenorhabditis elegans Innate Immune Response. Mol Cell Biol 27, 5544–5553.

Anderson, S. M., Cheesman, H. K., Peterson, N. D., Salisbury, J. E., Soukas, A. A. and Pukkila-Worley, R. (2019). The fatty acid oleate is required for innate immune activation and pathogen defense in Caenorhabditis elegans. Plos Pathog 15, e1007893.

Ayres, J. S. and Schneider, D. S. (2008). A Signaling Protease Required for Melanization in Drosophila Affects Resistance and Tolerance of Infections. Plos Biol 6, e305.

Behringer, D. C., Butler, M. J. and Shields, J. D. (2006). Avoidance of disease by social lobsters. Nature 441, 421–421.

Brenner, S. (1974). THE GENETICS OF CAENORHABDITIS ELEGANS. Genetics 77, 71– 94.

Brock, T. J., Browse, J. and Watts, J. L. (2006). Genetic Regulation of Unsaturated Fatty Acid Composition in C. elegans. Plos Genet 2, e108.

Brock, T. J., Browse, J. and Watts, J. L. (2007). Fatty Acid Desaturation and the Regulation of Adiposity in Caenorhabditis elegans. Genetics 176, 865–875.

Butcher, R. A., Ragains, J. R., Li, W., Ruvkun, G., Clardy, J. and Mak, H. Y. (2009). Biosynthesis of the Caenorhabditis elegans dauer pheromone. Proc. Natl. Acad. Sci. 106, 1875–1879.

Caldwell, R. M., Schafer, J. F., Compton, L. E. and Patterson, F. L. (1958). Tolerance to Cereal Leaf Rusts. Science 128, 714–715.

Carey, I. M., Critchley, J. A., DeWilde, S., Harris, T., Hosking, F. J. and Cook, D. G. (2018). Risk of Infection in Type 1 and Type 2 Diabetes Compared With the General Population: A Matched Cohort Study. Diabetes Care 41, 513–521.

Ciccarelli, E. J., Bendelstein, M., Yamamoto, K. K., Reich, H. and Savage-Dunn, C. (2024a). BMP signaling to pharyngeal muscle in the C. elegans response to a bacterial pathogen regulates anti-microbial peptide expression and pharyngeal pumping. Mol. Biol. Cell 35, ar52.

Ciccarelli, E. J., Wing, Z., Bendelstein, M., Johal, R. K., Singh, G., Monas, A. and Savage-Dunn, C. (2024b). TGF-β ligand cross-subfamily interactions in the response of Caenorhabditis elegans to a bacterial pathogen. PLOS Genet. 20, e1011324.

Clark, J. F., Meade, M., Ranepura, G., Hall, D. H. and Savage-Dunn, C. (2018). Caenorhabditis elegans DBL-1/BMP Regulates Lipid Accumulation via Interaction with Insulin Signaling. G3 Genes Genomes Genetics 8, 343–351.

Clark, J. F., Ciccarelli, E. J., Kayastha, P., Ranepura, G., Yamamoto, K. K., Hasan, M. S., Madaan, U., Meléndez, A. and Savage-Dunn, C. (2021). BMP pathway regulation of insulin signaling components promotes lipid storage in Caenorhabditis elegans. Plos Genet 17, e1009836.

Consortium*, T. C. elegans S. (1998). Genome Sequence of the Nematode C. elegans: A Platform for Investigating Biology. Science 282, 2012–2018.

Curtis, V. A. (2014). Infection-avoidance behaviour in humans and other animals. Trends Immunol 35, 457–464.

Dasgupta, M., Shashikanth, M., Gupta, A., Sandhu, A., De, A., Javed, S. and Singh, V. (2020). NHR-49 Transcription Factor Regulates Immunometabolic Response and Survival of Caenorhabditis elegans during Enterococcus faecalis Infection. Infect Immun 88, e00130–20.

Deshpande, R., Lee, B. and Grewal, S. S. (2022a). Enteric bacterial infection in Drosophila induces whole-body alterations in metabolic gene expression independently of the immune deficiency signaling pathway. G3 12, jkac163.

Deshpande, R., Lee, B., Qiao, Y. and Grewal, S. S. (2022b). TOR signaling is required for host lipid metabolic remodelling and survival following enteric infection in Drosophila. Dis Model Mech 15, dmm049551.

Desruisseaux, M. S., Nagajyothi, Trujillo, M. E., Tanowitz, H. B. and Scherer, P. E. (2007). Adipocyte, Adipose Tissue, and Infectious Disease. Infect Immun 75, 1066–1078.

Estevez, M., Attisano, L., Wrana, J. L., Albert, P. S., Massagué, J. and Riddle, D. L. (1993). The daf-4 gene encodes a bone morphogenetic protein receptor controlling C. elegans dauer larva development. Nature 365, 644–649.

Fanelli, M. J., Welsh, C. M., Lui, D. S., Smulan, L. J. and Walker, A. K. (2023). Immunity-linked genes are stimulated by a membrane stress pathway linked to Golgi function and the ARF-1 GTPase. Sci. Adv. 9, eadi5545.

Franz, A., Wood, W. and Martin, P. (2018). Fat Body Cells Are Motile and Actively Migrate to Wounds to Drive Repair and Prevent Infection. Dev Cell 44, 460–470.e3.

Gumienny, T. L. and Savage-Dunn, C. (2013). TGF-β signaling in C. elegans. Wormbook 1– 34.

Holdorf, A. D., Higgins, D. P., Hart, A. C., Boag, P. R., Pazour, G. J., Walhout, A. J. M. and Walker, A. K. (2020). WormCat: An Online Tool for Annotation and Visualization of Caenorhabditis elegans Genome-Scale Data. Genetics 214, 279–294.

Kavaliers, M., Choleris, E., Ågmo, A. and Pfaff, D. W. (2004). Olfactory-mediated parasite recognition and avoidance: linking genes to behavior. Horm Behav 46, 272–283.

Kim, D. H. and Ewbank, J. J. (2018). Signaling in the innate immune response. Wormbook 2018, 1–35.

Krishna, S., Maduzia, L. L. and Padgett, R. W. (1999). Specificity of TGFbeta signaling is conferred by distinct type I receptors and their associated SMAD proteins in Caenorhabditis elegans. Development 126, 251–260.

Laureano, G., Cavaco, A. R., Matos, A. R. and Figueiredo, A. (2021). Fatty Acid Desaturases: Uncovering Their Involvement in Grapevine Defence against Downy Mildew. Int J Mol Sci 22, 5473.

Li, S., Xu, S., Ma, Y., Wu, S., Feng, Y., Cui, Q., Chen, L., Zhou, S., Kong, Y., Zhang, X., et al. (2016). A Genetic Screen for Mutants with Supersized Lipid Droplets in Caenorhabditis elegans. G3 Genes Genomes Genetics 6, 2407–2419.

Liang, J., Yu, L., Yin, J. and Savage-Dunn, C. (2007). Transcriptional repressor and activator activities of SMA-9 contribute differentially to BMP-related signaling outputs. Dev Biol 305, 714–725.

Madhu, B., Lakdawala, M. F., Issac, N. G. and Gumienny, T. L. (2020). Caenorhabditis elegans saposin-like spp-9 is involved in specific innate immune responses. Genes Immun 21, 301–310.

Madhu, B., Lakdawala, M. F. and Gumienny, T. L. (2023). The DBL-1/TGF-β signaling pathway tailors behavioral and molecular host responses to a variety of bacteria in Caenorhabditis elegans. eLife 12, e75831.

Mallo, G. V., Kurz, C. L., Couillault, C., Pujol, N., Granjeaud, S., Kohara, Y. and Ewbank, J. J. (2002). Inducible Antibacterial Defense System in C. elegans. Curr Biol 12, 1209–1214.

Medzhitov, R., Schneider, D. S. and Soares, M. P. (2012). Disease Tolerance as a Defense Strategy. Science 335, 936–941.

Mochii, M., Yoshida, S., Morita, K., Kohara, Y. and Ueno, N. (1999). Identification of transforming growth factor-β-regulated genes in Caenorhabditis elegans by differential hybridization of arrayed cDNAs. Proc National Acad Sci 96, 15020–15025.

Morita, K., Chow, K. L. and Ueno, N. (1999). Regulation of body length and male tail ray pattern formation of Caenorhabditis elegans by a member of TGF-beta family. Development 126, 1337–1347.

Nandakumar, M. and Tan, M.-W. (2008). Gamma-Linolenic and Stearidonic Acids Are Required for Basal Immunity in Caenorhabditis elegans through Their Effects on p38 MAP Kinase Activity. PLoS Genet. 4, e1000273.

Nicholas, H. R. and Hodgkin, J. (2002). Innate Immunity: The Worm Fights Back. Curr Biol 12, R731–R732.

Olofsson, B. (2014). The olfactory neuron AWC promotes avoidance of normally palatable food following chronic dietary restriction. J. Exp. Biol. 217, 1790–1798.

O’Rourke, E. J., Soukas, A. A., Carr, C. E. and Ruvkun, G. (2009). C. elegans Major Fats Are Stored in Vesicles Distinct from Lysosome-Related Organelles. Cell Metab 10, 430–435.

Portal-Celhay, C., Bradley, E. R. and Blaser, M. J. (2012). Control of intestinal bacterial proliferation in regulation of lifespan in Caenorhabditis elegans. Bmc Microbiol 12, 49.

Råberg, L., Graham, A. L. and Read, A. F. (2009). Decomposing health: tolerance and resistance to parasites in animals. Philosophical Transactions Royal Soc B Biological Sci 364, 37–49.

Read, A. F., Graham, A. L. and Råberg, L. (2008). Animal Defenses against Infectious Agents: Is Damage Control More Important Than Pathogen Control? Plos Biol 6, 10.1371/journal.pbio.1000004.

Robertis, E. M. D. (2008). Evo-Devo: Variations on Ancestral Themes. Cell 132, 185–195.

Roberts, A. F., Gumienny, T. L., Gleason, R. J., Wang, H. and Padgett, R. W. (2010). Regulation of genes affecting body size and innate immunity by the DBL-1/BMP-like pathway in Caenorhabditis elegans. Bmc Dev Biol 10, 61.

Savage, C., Das, P., Finelli, A. L., Townsend, S. R., Sun, C. Y., Baird, S. E. and Padgett, R. W. (1996). Caenorhabditis elegans genes sma-2, sma-3, and sma-4 define a conserved family of transforming growth factor beta pathway components. Proc National Acad Sci 93, 790– 794.

Schafer, J. F. (1971). Tolerance to Plant Disease. Annu Rev Phytopathol 9, 235–252.

So, S., Tokumaru, T., Miyahara, K. and Ohshima, Y. (2011). Control of lifespan by food bacteria, nutrient limitation and pathogenicity of food in C. elegans. Mech Ageing Dev 132, 210–212.

Sternberg, P. W., Auken, K. V., Wang, Q., Wright, A., Yook, K., Zarowiecki, M., Arnaboldi, V., Becerra, A., Brown, S., Cain, S., et al. (2024). WormBase 2024: status and transitioning to Alliance infrastructure. GENETICS 227, iyae050.

Suzuki, Y., Yandell, M. D., Roy, P. J., Krishna, S., Savage-Dunn, C., Ross, R. M., Padgett, R. W. and Wood, W. B. (1999). A BMP homolog acts as a dose-dependent regulator of body size and male tail patterning in Caenorhabditis elegans. Development 126, 241–250.

Tenor, J. L. and Aballay, A. (2008). A conserved Toll like receptor is required for Caenorhabditis elegans innate immunity. Embo Rep 9, 103–109.

Urist, M. R. and Strates, B. S. (1971). Bone Morphogenetic Protein. J Dent Res 50, 1392– 1406.

Watts, J. L. and Browse, J. (2002). Genetic dissection of polyunsaturated fatty acid synthesis in Caenorhabditis elegans. Proc. Natl. Acad. Sci. 99, 5854–5859.

Yamamoto, K. K. and Savage-Dunn, C. (2023). TGF-β pathways in aging and immunity: lessons from Caenorhabditis elegans. Front. Genet. 14, 1220068.

Yu, Y., Mutlu, A. S., Liu, H. and Wang, M. C. (2017). High-throughput screens using photo-highlighting discover BMP signaling in mitochondrial lipid oxidation. Nat Commun 8, 865.

Zhang, S. O., Box, A. C., Xu, N., Men, J. L., Yu, J., Guo, F., Trimble, R. and Mak, H. Y. (2010). Genetic and dietary regulation of lipid droplet expansion in Caenorhabditis elegans. Proc National Acad Sci 107, 4640–4645.

Zugasti, O. and Ewbank, J. J. (2009). Neuroimmune regulation of antimicrobial peptide expression by a noncanonical TGF-β signaling pathway in Caenorhabditis elegans epidermis. Nat Immunol 10, 249–256.

